# How woodcocks produce the most brilliant white plumage patches among the birds

**DOI:** 10.1101/2022.12.09.519795

**Authors:** Jamie Dunning, Anvay Patil, Liliana D’Alba, Alexander L Bond, Gerben Debruyn, Ali Dhinojwala, Matthew Shawkey, Lukas Jenni

## Abstract

Until recently, and when compared with diurnal birds that use contrasting plumage patches and complex feather structures to convey visual information, communication in nocturnal species was considered to follow acoustic and chemical channels. However, many nocturnal birds have evolved intensely white plumage patches within otherwise inconspicuous plumages. We used spectrophotometry, electron microscopy, and optical modelling to explain the mechanisms producing bright white tail feather tips of the Eurasian woodcock *Scolopax rusticola*. Their diffuse reflectance was ∼30% higher than any previously measured feather. This intense reflectance is the result of incoherent light scattering from a disordered nanostructure composed of keratin and air within the barb rami. In addition, the flattening, thickening, and arrangement of those barbs creates a Venetian-blind-like macrostructure that enhances the surface area for light reflection. We suggest that the woodcocks have evolved these bright white feather patches for long-range visual communication in dimly lit environments.

## 1. Introduction

The use of contrasting plumage patches or complex feather structures to convey information is widespread in birds (reviewed in Jenni and Winkler 2020; Terrill and Shultz 2022). Unlike in diurnal birds, visual signals in nocturnal species are understudied, and communication was, until recently, considered to follow chemical and acoustic channels (Healy and Guilford 1990; Bonadonna and Bretagnolle 2002; Grieves et al. 2022). However, in dim light environments, plumage characteristics have emerged that maximize reflectance of available light (Endler 1993; Penteriani and Del Mar Delgado 2017). While most nocturnal birds have inconspicuous or cryptic plumages, visual signals are typically intensely white; for example, the white patches in the plumage of some nightjars Caprimulgidae (Aragonés, Arias De Reyna, and Recuerda 1999), true owls Strigidae (Penteriani et al. 2007; Bortolotti, Stoffel, and Galván 2011; Bettega et al. 2013), stone-curlews Burhinidae (Cramp and Simmons 1983), and snipes Scolopacidae (Höglund, Eriksson, and Lindell 1990).

The function and the mechanism by which these white patches optimise light reflectance is not well understood (but see Igic, D’Alba and Shawkey 2016; Igic, D’Alba and Shawkey 2018), but they communicate behavioural intention, for example, mating or territorial behaviours, or signal quality (Höglund, Eriksson, and Lindell 1990; but also see Sæther et al. 2000). However nocturnal birds typically also require crypsis while roosting during day light (Troscianko et al. 2016; Stevens et al. 2017) and therefore conceal their visual signals. White wing patches of some nightjars are, for example, only exposed in flight (Aragonés, Arias De Reyna, and Recuerda 1999), or, in the woodcocks *Scolopax* spp, white undertail feather patches are only exposed when the tail is raised (Borodulina and Formosow 1967; Figure 2.Ca - b).

Borodulina and Formosow (1967) first described modifications to the rami that radiate from the central rachis of the feather) that comprise the white tips on the underside of the Eurasian woodcock’s *Scolopax rusticola* (hereafter woodcock) tail feathers (hereafter rectrices) but did not measure reflectance and characterise its mechanism. Previous studies have demonstrated how micro-structures correlate with white plumage intensity, for example in the winter body plumage of the rock ptarmigan *Lagopus muta* (Dyck 1979), the opal-like colours on some manakin birds Pipridae (Igic, D’Alba, and Shawkey, 2016) and between many white-plumaged birds from different families (Igic, D’Alba, and Shawkey, 2018). Likewise, ‘super-white’, derived of micro-structures on the carapace of a beetle (Vukusic et al., 2007; Burresi et al., 2014) were well reported. The white patches in nocturnal birds, which are potentially optimised for signalling in low-light conditions, have seldom been addressed and require more detailed analysis.

Here we describe the mechanisms by which the white rectrix tips of the woodcock produce an intense white signal in low light conditions, using angle-resolved and diffuse spectrophotometry, electron microscopy and optical modelling via finite-difference time-domain (FDTD) approaches.

## 2. Material and Methods

### (a) Microscopy

To characterize the microstructure and nanostructure responsible for producing the bright white signal, we used scanning and transmission electron microscopy (SEM and TEM, respectively). For SEM, we mounted individual white and brown rami (obtained from the same feather) separately, on stubs with carbon tape. We also oriented small fragments of rami in a way that allowed their observation in cross section. We sputter-coated the samples with gold/palladium for 2 minutes and imaged them on a SEM (FlexSEM 1000; Hitachi) at an accelerating voltage of 10 kV and 6 mm working distance.

For TEM we first embedded individual rami following a standard protocol (D’Alba et al 2021). Briefly, we rinsed and dehydrated the rami using ethanol three times, and then infiltrated them with increasing concentrations (15%, 50%, 70% and 100%) of epoxy resin (EMbed-812; Electron Microscopy Sciences, PA, USA) followed by 16-hour polymerization in epoxy resin at 60° C in a laboratory oven.

We trimmed the blocks containing the rami and cut 100 nm thick cross sections using a Leica UC-6 ultramicrotome (Leica Microsystems, Germany). We collected the sections using oval-slit carbon and formvar-coated copper grids in duplicate and stained with Uranyless/lead citrate. We observed the sections on a JEOL JEM 1010 (Jeol Ltd, Tokyo, Japan) transmission electron microscope operating at 120 kV.

### (b) Spectrophotometry

We used micro- and (macro)spectrophotometry to measure light reflectance from three separate rectrices. We measured reflectance from the reverse surface of a white ramus using a micro-spectrophotometer (CRAIC AX10: sensitivity 320-800 nm); and a spectrophotometer that measured a region across several rami (∼2 mm spot size). We measured diffuse (all reflected light) and specular reflectance (light reflected at a specific angle) between 300 - 700 nm in increments of 1 nm using a AvaSpec-2048 spectrometer and dual light source set-up (AvaLight-DH-S deuterium-halogen light source and AvaLight-HAL-S-MINI light source). We measured diffuse reflectance (which assumes that light reflectance is influenced by internal structures as well as those on an object’s surface) using a bifurcated probe and an integrating sphere with a black gloss trap to exclude specular (light reflected from an objects surface) reflectance (AvaSphere-50-REFL). Then, we measured specular reflectance at three different angles (75°, 60°, 45°) using a bifurcated probe and a block holder (AFH-15, Avantes). We placed each feather on black paper minimizing background reflectance. All measurements are expressed relative to an 99% white reflectance standard (WS-2, Avantes) and 2% Avantes black standard (BS-2, Avantes). We processed data in the R package pavo in R 4.1.2 (Maia et al. 2019; R Core Team 2022) and plotted them with previously published measurements from 61 other birds using identical spectrophotometric methods (Igic, D’Alba and Shawkey 2018).

### (c) Finite-Difference Time-Domain (FDTD) simulations

To explore the directionality of reflectance as a function of varying rami angle, we modelled how photons interact with structures within an individual barb. We ran a series of finite-difference time-domain (FDTD) simulations using a commercial-grade Ansys Lumerical 2021 R1 solver (Ansys, Inc.). The FDTD method provides a general solution to any light scattering problem on complex arbitrary geometries (in this case, a unit cell structure of an individual ramus) by numerically solving Maxwell’s curl equations on a discrete spatiotemporal grid (Taflove and Hagness 2005). The simulation estimates all scattered light at all angles and, in this respect, is not directly comparable with our diffuse spectrophotometry data.

Our simulated 3D CAD models were based on empirical microscopic observations of the woodcock barbs (see supplementary material, S1:A-D). First, we rendered a 3D CAD geometry for a control hollow unit cell, without internal photonic nanostructures, and a solid unit cell. We used SEM microscopy to define CAD dimensions, each cell had a keratin cortex thickness of 7 μm with a hollow interior, 20 μm high (Z direction) and 8 μm wide (X direction). We then used SEM microscopy to render a unit cell with an internal nanostructure equivalent to the woodcock’s rami, i.e. of air pockets and a supporting matrix of nano-fibres (see Figure 1). We did this using a uniform random distribution of non-overlapping spherical particles within the keratin matrix, which randomly varied in diameter between 0.45 μm and 3.45 μm. The optical constants (complex refractive indcies) for keratin were adapted from previous literature (Stavenga et al. 2015; Table S1).

**Figure 1.**
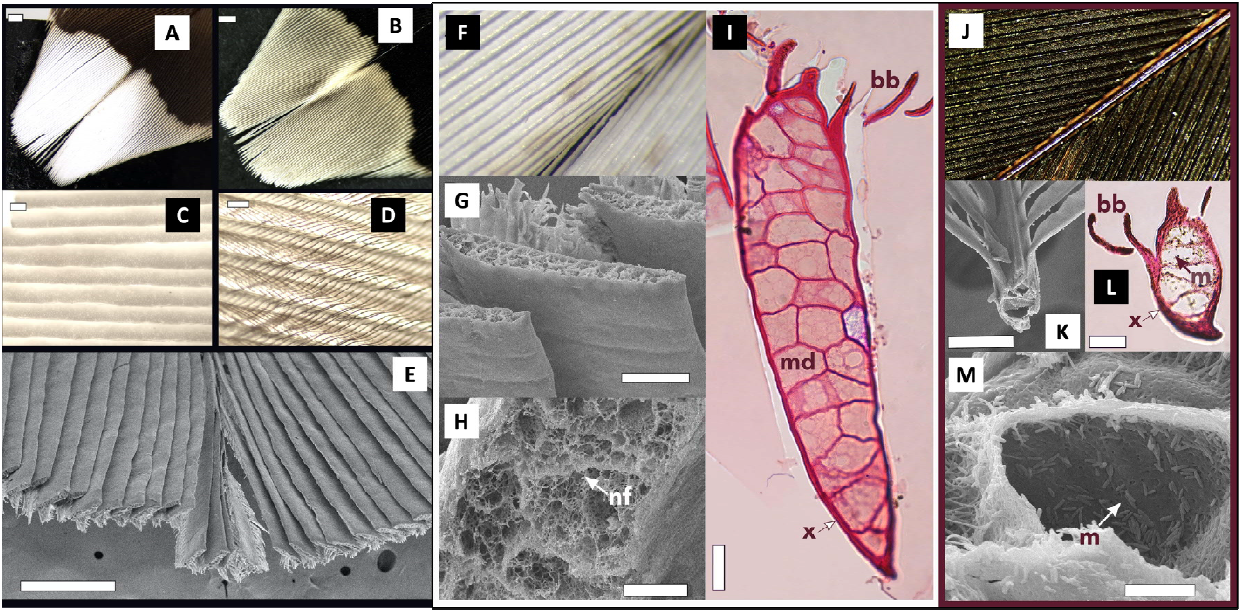
**A) – E): Morphology of the white tips of woodcock *Scolopax rusticola* rectrices.** A) White reverse surface. B) Brown obverse surface. C) White rami in a Venetian-blind alignment; individual cells are apparent. D) Obverse view showing the interlocked dark barbules covering the white rami. E) SEM micrograph of the white rectrix tip transversally cut, showing shallow V-shaped surface of rami; **F) – M): Comparison of the microstructure of the white and brown parts of rectrices**. F) Optical image of white rami. G) Thickened and flattened rami viewed from the reverse surface. H) Interior of a white ramus shows cells with networks of keratin fibres (nf) and air pockets. I) a white ramus showing hollow medullary cells (md) and a thin cortex (x); the barbules (bb) are present on the obverse side. J) Optical image of contiguous brown region. K) Brown rami in cross-section. L) Melanosomes (m) present throughout the rami and barbules. M) Medullary cell of brown ramus showing melanosomes (m) and the absence of keratin matrices. Scale bars: A and B) 1mm; C), D), G) and K) 50μm; E) 500μm; H) 10μm; I) and L) 100μm; M) 5μm.

We performed simulations using a broadband plane wave source (400-700 nm), propagated along the -Z direction. First, at a normal angle of incidence (AOI; 0° from cell surface) and then at 70°, for our control, hollow and solid, unit cells. Then, we ran simulations using our simulated woodcock cell at 0°, 20°, 50°, 70° and 80° AOI. Boundary conditions in the lateral direction (X and Y) were set to periodic. We monitored reflectance data using a Discrete Fourier Transform (DFT) power monitor placed behind the source injection plane. The simulation time (in fs) and boundary condition along the light propagation direction (Z; perfectly matching layer (PML) boundaries) were chosen such that the electric field decayed before the end of the simulation (auto-shutoff criteria). All the incident light was either reflected, transmitted, or absorbed.

## 3. Results

### (a) Structure of the white rectrix tips

The tips of the rectrices are white on the reverse (figures 1A and 2A), but greyish brown on the obverse surface (figure 1B). The rami are thickened and flattened in the white patch and overlap each other, superficially like Venetian-blinds (figures 1C, 1E). The angle of these rami relative to the feather surface vary (as suggested by Borodulina and Formosow 1967), we estimated from ∼70º for proximal rami to ∼76º for distal rami (figure 1E). The proximal and distal brown barbules originate from the upper surface of the rami, hence are only visible on the obverse surface and cover the thickened white rami from above, providing the greyish brown colour of the obverse surface (figures 1B and 1D). They interlock to form a coherent vane. The two sides of a white tip, separated by the rachis, are concave and the barbs arranged in opposite angles (figure 1J), reflecting light in different directions and apparent when turning a feather in low light. In contrast, the brown parts of the rectrices are structurally typical of vaned feathers with thin barbs that are spaced by the brown barbules (figure 1J, 1K). The thickened white rami in the feather tips were ∼2.5 times thicker and appeared internally more complex than brown rami (figure 1 F-H and 1J-M, respectively). The medulla of white rami contained numerous and complex photonic cells with fine networks of nanofibers and scattered air pockets (figure 1G-I), lacking melanosomes entirely. These matrices of air and keratin appeared disorganized. In contrast, rami from brown feather regions were less thick, rounder, had fewer medullary cells and did not contain a matrix of air and keratin, but were abundant in melanosomes both inside the barb medulla and the cortex (figure 1K-M).

**Figure 2.**
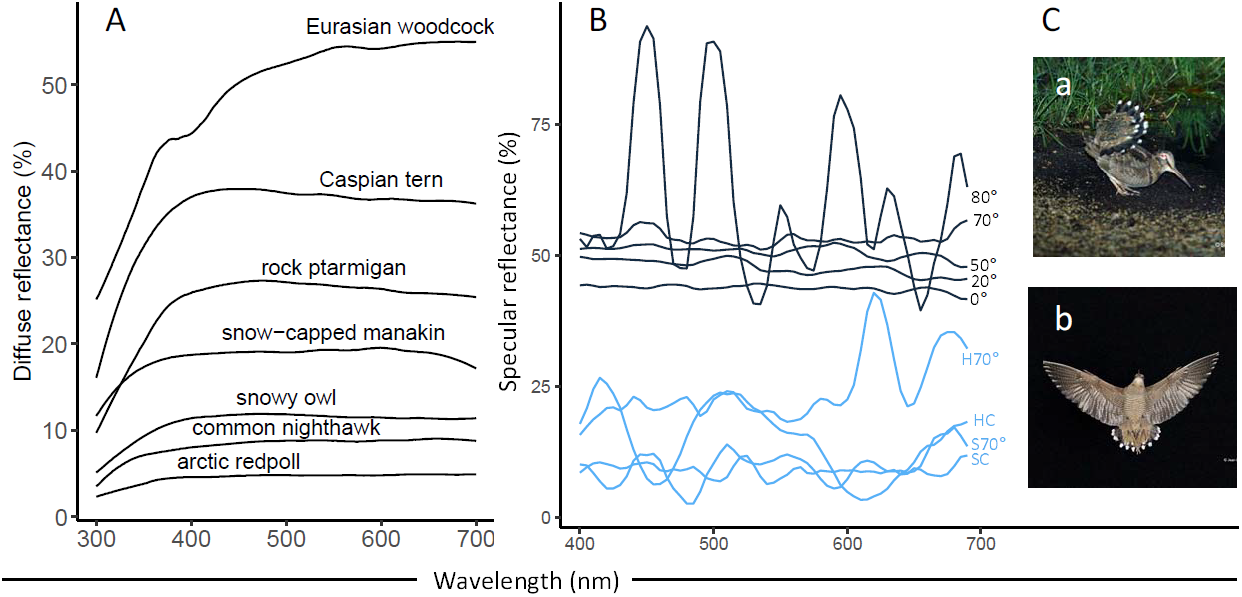
(A) Diffuse reflectance spectra measured from the reverse surface of the white Eurasian woodcock *Scolopax rusticola* rectrix tips, peaking at ∼55%, 31% brighter than the next brightest feather, Caspian tern *Hydroprogne caspia*, and compared against 61 white plumages from Igic, D’Alba, and Shawkey (2018), species mentioned in text are highlighted; (B) Finite-Difference Time-Domain (FDTD) simulations showing simulated reflectance at five angles of incidence (AOI; highlighted in grey, 0, 20, 50, 70 and 80) and four control measurements (highlighted in blue; hollow cell at 70 AOI, hollow cell control, solid cell at 70 AOI and solid control). These data suggest that air pockets present in the keratin matrix are essential for increasing the reflectivity across visible wavelengths in the woodcock’s tail feathers. (C) Showing ecological context when white tips are exposed, either from the ground (probably a female attracting an overflying male) (Ca) or in flight (male in display flight) (Cb); photos by Serge Santiago and Jean-Lou Zimmermann,

### (b) Reflectivity

Spectrophotometry revealed intense diffuse reflectance across rami on the white underside of the rectrices, peaking at 55% (628 nm) (figure 1F; 2A). Likewise, individual rami had even greater specular reflectance, peaking >100% against a diffuse standard (supplementary figure S2). The white patches on woodcock rectrices are therefore exceptionally bright, and, to the best of our knowledge, represent the brightest white measured from the plumage of a bird, 31% brighter than the next most reflective, Caspian tern *Hydroprogne caspia*, that peaks at 38% (459nm), and 91% brighter than the least-reflective white feather measured, arctic redpoll *Acanthis hornemanni*, that peaks at 4.9% (638nm) (Igic, D’Alba, and Shawkey 2018; figure 2A). Specular reflectance was highest when measured at 75º relative to surface normal, decreasing at more acute angles, suggesting some directionality to reflectance intensity (supplementary figure S3).

### (c) Finite-Difference Time-Domain simulations of reflectivity

We found the disordered nanostructure formed by keratin and air phases in the woodcock rami were essential for generating intense white reflectance. For normal incidence (0° from the surface normal), the overall reflectance for the woodcock-mimicked rami unit cell nanostructure increased by ∼65% with respect to the control hollow unit cell nanostructure. Additionally, the simulations also highlight some directionality to patch intensity. Modelled reflectance at 80°, although showed high reflectance, also showed increased noise, which we suggest is due to interference effects on the surface of the feather structures. Otherwise, the reflectance increased from a peak of ∼45% at normal incidence (0°), to a peak of ∼57% at 70°, which represents the actual angle of the rami within the white patch (figure 2B). Reflectance at 70° is broadly the same as the actual diffuse reflectance (figure 2A), although FDTD simulates diffuse plus specular reflectance. We therefore suggest that the rami are arranged to lie at the angle which best optimizes reflectance. Further, our simulated control cells demonstrate that air pockets in the keratin matrix are essential for increasing the overall reflectivity across visible wavelengths.

## Discussion

Our results suggest that the white tips on the woodcock’s rectrices represent the brightest reflectance yet measured and, by virtue, the whitest white plumage patch currently known among the birds. Other bright white plumages have been reported previously, but they are either supposition (Tickell 2003), or using different methods or without standardised comparison (Dyck 1979; Caswell & Prum 2011). We present our results alongside those previously described plumages (see Igic, D’Alba, and Shawkey 2018 for a full list), using standardised a approach (Figure 2A). This reflectance is produced by the arrangement of thick and flattened rami with a broad distribution of air pockets, that together maximize light reflectance. We used FDTD simulations to demonstrate that 1) the internal structure of the rami on the white tips is integral for light scattering and subsequent reflectance, but also; 2) that the angle of the broadened barbs in relation to each other optimise reflectance at the macro-scale.

The structures we describe differ from those of less intense diurnal plumages in two ways: First, the rami are thickened and flattened (Borodulina and Formosow 1967; this study), increasing surface area available for reflection and preventing light from passing between the rami and barbules. Second, the thickened rami allow for a complexity of photonic cells, with a network of keratin nanofibers and scattered air pockets, creating numerous interfaces to favour scattering events (which similar to the ‘super-white’ reflectance described in a white beetle; Vukusic 2007, Burresi et al. 2014).

Igic, D’Alba, and Shawkey (2018) suggested that more intense reflectance of white plumage was associated with densely packed, rounder and less hollow rami, but also thicker and longer barbules. Consequently, larger species were brighter by virtue of rami thickness and complexity. However, the woodcock rami are thickened and flattened, superficially like the rami in the white crown of Blue-rumped Manakin *Lepidothrix isidorei* (Igic, D’Alba, and Shawkey 2016); in this case, the internal nanostructure is without the thickened rami that increases the surface area of reflectance. Despite some similarities, the diffuse reflectance of the manakin’s crown peaks at ∼17% (Igic, D’Alba, and Shawkey 2016), ∼105% less bright than the woodcock. However, specular reflectance of the manakin crown is higher than the woodcock, due to a nanostructure that enhances specular reflectance (also see Shawkey, Maia and D’Alba 2011; McCoy et al. 2021).

The Venetian-blind arrangement of the thickened rami, and subsequent directional reflectance, is like the arrangement of barbules of hummingbirds Trochilidae. Here, the angle of the barbules relative to the axis of the ramus, and the angle between the proximal and distal barbules of the rami determine directionality of reflectance, associated with irradiance (Giraldo, Sosa and Stavenga, 2021).

White patches are present in all eight species of woodcock, but not in their closest relatives (23 species of non-*Scolopax* Scolopacidae, see supplementary table S1) and signal some behavioural intention in dimly lit environments (Cramp and Simmons 1983; Glutz von Blotzheim et al., 1977). Because these patches are only visible from below, any functional significance is conditional on raising and fanning the tail, for example during courtship displays (Hagen, 1950; Hirons, 1980; Ferrand and Gossmann 2009; Lastukhin and Isakov, 2016), predator distraction or non-reproductive communication (Ingram, 1974; Fetisov, 2017). The link between patch intensity, behaviour and relative light environment is understudied and would benefit from further research.

We suggest that the woodcocks have evolved brilliant white feather patches, the brightest described within the birds, through elaborate structural modifications at the macro-, micro- and nano scales for communication in dimly lit environments.

## Supporting information

supp. matt.

## Acknowledgments

We would like to thank Mary Hennen, Michael A Pearson, Matt Rayner, Paul Sweet, Arseny Tsvey, Niklaus Zbinden for conversation, samples, and translations, and Serge Santiago and Jean-Lou Zimmermann for the photos. AD and AP acknowledge financial support from the US Air Force Office of Scientific Research (AFOSR) under Multidisciplinary University Research Initiative (MURI) grant (FA 9550-18-1-0142).

## Ethics statement

We sourced woodcock rectrices from a private collection from Switzerland without the need for specific licensing.

