## Supplementary material for "How woodcocks produce the most brilliant white plumage patches among the birds": supp. matt.

Figure S1: Schematic representation of the 3D CAD geometries used in the Finite-Difference Time-Domain simulations of reflectivity. The schematics represent the XZ view of the 3D geometry. The downward pink arrow indicates the direction of propagation of light through the system and the horizontal blue arrow indicates the polarization state of incident light (in this case, *p*-polarization state).

(A) Hollow barb unit cell structure (angle of incidence (AOI) 0 degrees).

(B) Simulated Eurasian Woodcock white feather barb unit cell structure (AOI 0 degrees)

(C) Simulated Eurasian Woodcock white feather barb unit cell structure (AOI 20 degrees)

(D) Simulated Eurasian Woodcock white feather barb unit cell structure (AOI 50 degrees)

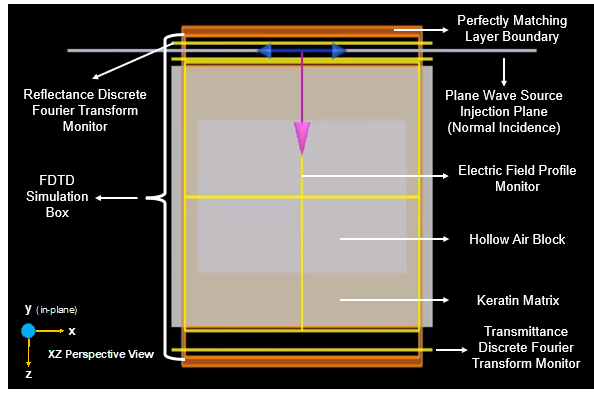
(D) Simulated Eurasian Woodcock white feather barb unit cell structure (AOI 70 degrees)

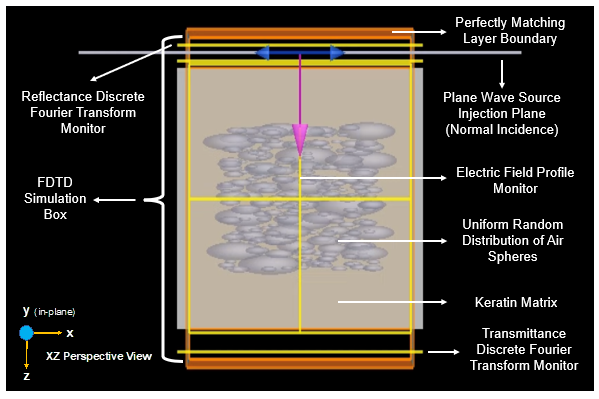

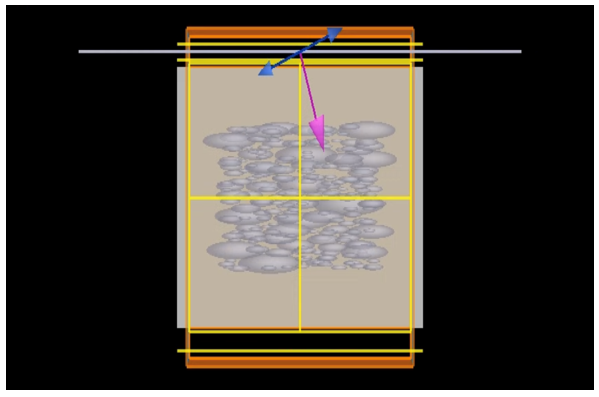

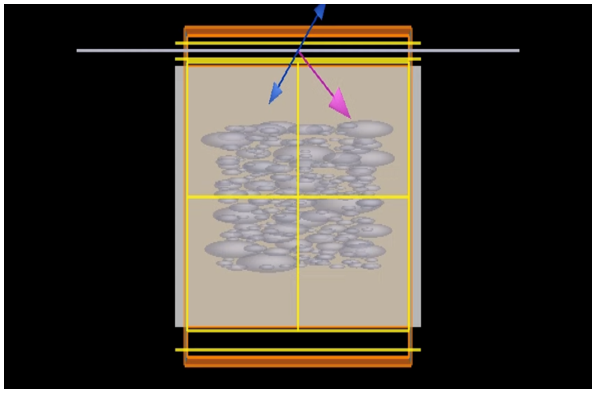

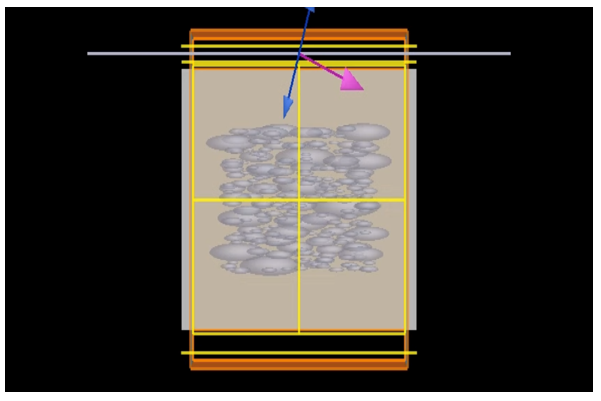

Figure S2-S3. Specular reflectance was greatest in the white tail feather tips, compared with the grey obverse surface of the tail feather tip, and the orange and brown barring on the obverse surface of the feather away from the tip, and at ~75° (S3), suggesting some directionality to patch intensity.

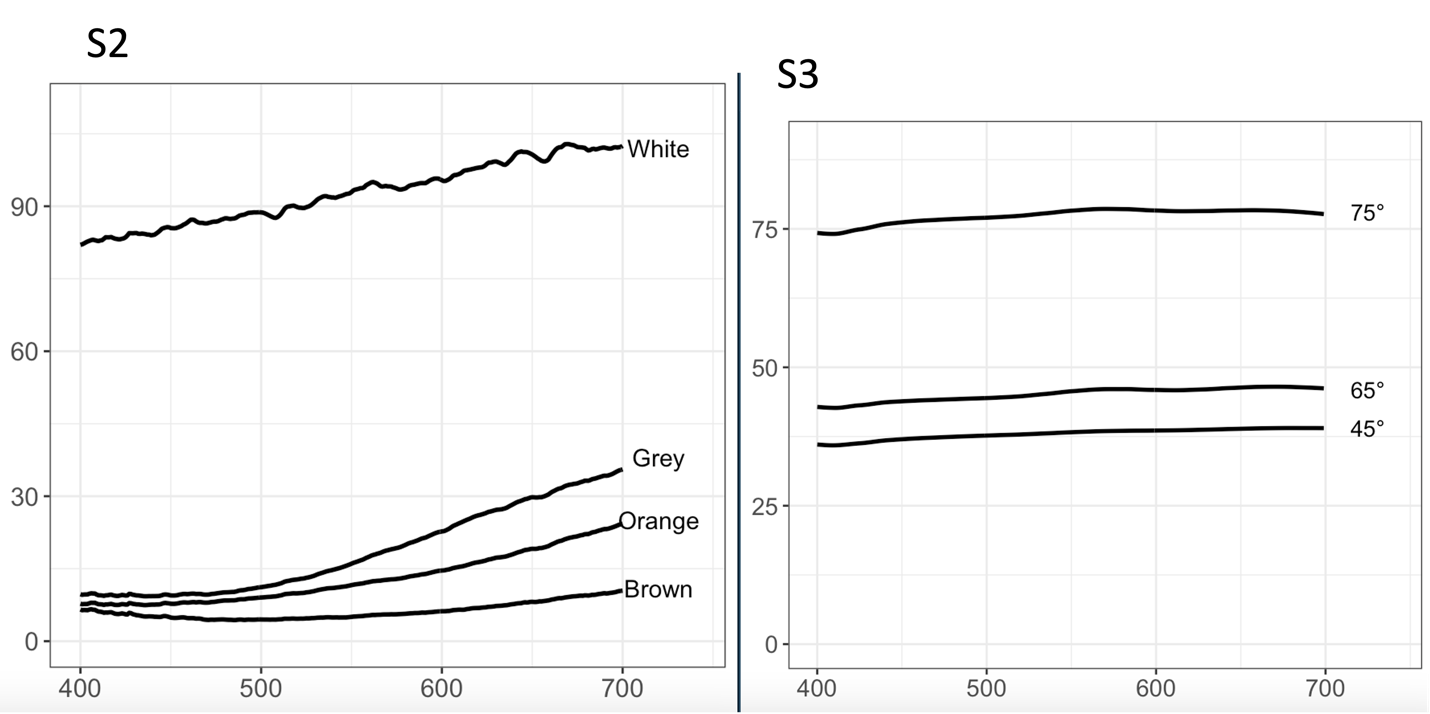

Table S1. Museum specimens were accessed from four institutions to check for broadened white rami in the tail: Natural History Museum, Tring, UK (NHM Tring); the Field Museum Natural History, Chicago, USA (FMNH); American Museum of Natural History, New York, USA (AMNH); and, the Auckland Museum, New Zealand (AMNZ).

| Species | Common name | Collection |
| --- | --- | --- |
| *rusticola* | Eurasian woodcock | This study |
| *minor* | American woodcock | NHM Tring |
| *mira* | Amani woodcock | NHM Tring |
| *bukidnonensis* | Bukidnon woodcock | FMNH |
| *saturata* | Javan woodcock | NHM Tring |
| *rosenbergii* | New Guinea woodcock | NHM Tring |
| *rochussenii* | Moluccan woodcock | NHM Tring |
| *celebensis* | Sulawesi woodcock | AMNH |
| *minimus* | Jack snipe | NHM Tring |
| *megala* | Swinhoe’s snipe | NHM Tring |
| *stenura* | Pin-tailed snipe | NHM Tring |
| *media* | Great snipe | NHM Tring |
| *undulata* | Giant snipe | NHM Tring |
| *jamesoni* | Jameson’s snipe | NHM Tring |
| *stricklandii* | Fuegian snipe | NHM Tring |
| *hardwickii* | Latham’s snipe | NHM Tring |
| *nemoricola* | Wood snipe | NHM Tring |
| *macrodactyla* | Madagascar snipe | NHM Tring |
| *nobilis* | Noble snipe | NHM Tring |
| *andina* | Puna snipe | NHM Tring |
| *paraguaiae* | Pantanal snipe | NHM Tring |
| *gallinago* | Common snipe | NHM Tring |
| *nigripennis* | African snipe | NHM Tring |
| *soliitaria* | Solitary snipe | NHM Tring |
| *Imperialis* | Imperial snipe | NHM Tring |
| *delicata* | Wilson’s snipe | NHM Tring |
| *barrierensis* | North Island snipe | AMNZ |
| *iredalei* | South Island snipe | NHM Tring |
| *pusilla* | Chatham island snipe | NHM Tring |
| *huegeli* | Snares island snipe | NHM Tring |
| *aucklandica* | Subantarctic snipe | NHM Tring |
